# A non-zero variance of Tajima’s estimator for two sequences even for infinitely many unlinked loci

**DOI:** 10.1101/069989

**Authors:** Léandra King, John Wakeley, Shai Carmi

**Affiliations:** Department of Organismic and Evolutionary Biology, Harvard University, Cambridge, MA, USA; Braun School of Public Health and Community Medicine, The Hebrew University of Jerusalem, Israel

**Keywords:** Coalescent Theory, Recombination, Heterozygosity, Effective Population Size, Pedigrees, Genealogies, Markov Chains

## Abstract

The population-scaled mutation rate, *θ*, is informative on the effective population size and is thus widely used in population genetics. We show that for two sequences and *n* unlinked loci, Tajima’s estimator (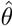), which is the average number of pairwise differences, is not consistent and therefore its variance does not vanish even as *n* → ∞. The non-zero variance of 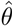 results from a (weak) correlation between coalescence times even at unlinked loci, which, in turn, is due to the underlying fixed pedigree shared by all genealogies. We derive the correlation coefficient under a diploid, discrete-time, Wright-Fisher model, and we also derive a simple, closed-form lower bound. We also obtain empirical estimates of the correlation of coalescence times under demographic models inspired by large-scale human genealogies. While the effect we de scribe is small 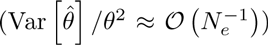, it is important to recognize this feature of statistical population genetics, which runs counter to commonly held notions about unlinked loci.

## 1 Introduction

The population mutation rate, *θ*, is defined as 4*N*_e*μ*_, where *N*_e_ is the effective population size and *μ* is the mutation rate per locus per generation. Two classic estimators were developed for *θ*, Watterson’s (based on the number of segregating sites (Watterson, 1975)) and Tajima’s (based on the average number of pairwise differences (Tajima, 1983, 1989)). For a single pair of sequences, both estimators are identical (denoted here as 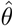) and equal to the number of differences between the sequences.

Increasing the number of sampled individuals has limited ability to improve these estimates of *θ*, because shared ancestry reduces the number of independent branches on which mutations can arise (Rosenberg and Nordborg, 2002). Felsenstein (2006) showed that the variance of maximum likelihood estimates of *θ* decreases approximately logarithmically with the number of individuals sampled. In contrast, the variance decreases inversely with the number of independent loci. Thus, to increase the accuracy of estimates of *θ*, it is generally more effective to increase the number of independent loci than the sample size at each locus (see also e.g., (Pluzhnikov and Donnelly, 1996) and references within).

Consider a set of *n* unlinked loci located on different (non-homologous) chromosomes. We show here that even as *n* → ∞, the variance of the resulting estimate of *θ* does not converge to zero, in contrast to what we may have naïvely assumed. This behavior results from the fact that coalescence times, even at unlinked loci, are in fact weakly correlated, due to the sharing the same fixed underlying pedigree across all genealogies at all loci (Wakeley et al., 2012). By conditioning on the number of shared genealogical common ancestors, we derive a simple lower bound, as a function of *N*_*e*_, on the variance of 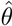.

Unlinked loci may also be sampled from the same chromosome, separated by an infinitely high recombination rate. The correlation of coalescence times in such a case is higher, as the two loci may travel together for the first few generations. Therefore, the extent of the correlation, and thereby, the variance of 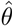, also depend on the *sampling configuration*. We derive the correlation coefficient analytically, as a function of the configuration and the effective population size, using a diploid discrete time Wright-Fisher model (DDTWF). This model is an extension of the haploid DTWF model, previously advocated by Bhaskar et al. (2014) for the study of large samples from finite populations.

Our results for the variance of 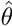 were obtained under the Wright-Fisher demographic model. To shed light on the variance of 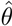 under more realistic demographic models, we run simulations based on real, large-scale human genealogical data (Erlich, 2016). The pedigrees inspired by different human populations differ from each other and from the Wright Fisher pedigrees in a number of ways, for example in the variance of the relatedness of any two randomly chosen individuals. These differences lead to differences in the variance of 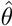 for each population, even if they have the same effective population size.

## 2 The relation of the variance of 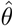 to the correlation of the coalescence times

For a sample of size two at *n* loci, the estimator of *θ* can be expressed as

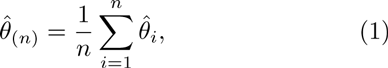

where 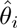 is the number of differences at locus *i*. If we assume the loci are exchangeable, we have:

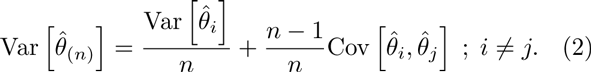

Under the standard coalescent model (Kingman, 1982), 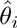 is Poisson distributed with mean 2*μT_i_*, where *T_i_* is the time until coalescence at locus *i* in generations and *μ* is the mutation rate per locus per generation. Using the law of total covariance,

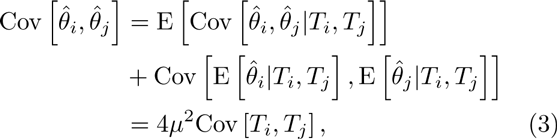

since conditional on *T_i_* and *T_j_*, 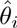 and 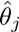 are independent. Thus, for infinitely many sites,

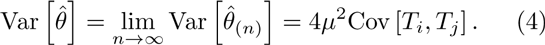

Because *T_i_* is distributed exponentially with rate 1/(2*N_e_*) under the standard coalescent model (Kingman, 1982; Tajima, 1983), 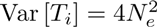. Since Cov [*T*_*i*_,*T*_*j*_] = Corr [*T*_*i*_,*T*_*j*_] × Var [*T*_*i*_], we can write:

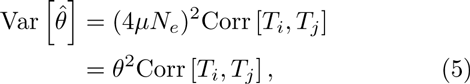

and we focus henceforth on the correlation of *T*_*i*_ and *T*_*j*_. Studying the correlation instead of the covariance allows us, later on, to visually compare the results across different effective population sizes.

We note that the variance of 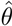 is calculated over independent repeats over the entire evolutionary process, including the generation of the population pedigree (family relationships between all individuals), as well as the gene genealogies. We elaborate below on this important point (sections 3, 5, and the Discussion).

## 3 Modeling the effect of the shared pedigree

In this section, we provide an intuitive derivation of the role of the shared underlying pedigree in generating a non-zero variance of 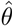.

### 3.1 Inconsistency of 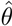 due to the underlying pedigree

We begin with a general analysis of the inconsistency of the estimator of *θ*. The value of 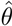 is a function of the pedigree that connects the two individuals in our sample, where the pedigree itself is randomly drawn from a demographic model (e.g., the Wright-Fisher model) with parameter *θ*. If the sampled individuals happen to be more closely related than average, then 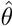 will tend to underestimate the true value of *θ*. The opposite is true if the sampled individuals are less closely related than average.

Let *δ* be the probability that a randomly sampled pair of individuals is very closely related, for example as full siblings. Let *ϵ* be some arbitrary value smaller than the difference between *θ* and 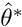, where 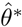 is estimated from a sample of full siblings. By sampling sufficiently many loci (or gene genealogies), we could theoretically infer the common ancestry of the sampled pair to any desired accuracy. However, this would not give information about the pedigree beyond the ancestry of the sampled pair, and as the sampled pair is related more closely than average, 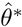 would underestimate *θ*. For this fixed *ϵ* and *δ*, we therefore cannot find n large enough such that 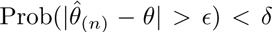. This implies that is no convergence in probability, which means that this estimate of *θ* is not consistent. In turn, this inconsistency implies that the variance of 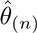 does not tend to 0 as *n* increases.

### 3.2 A lower bound on the limiting variance

Next, we derive an intuitive lower bound on the limiting variance of 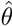 for a sample of two loci on non-homologous chromosomes, where according to Eq. (4), we only need the covariances of *T*_*i*_ and *T*_*j*_. To compute these covariances, we condition on a vector of variables {*x*} = *x*_1_, *x*_2_,…, *x_G_*, where *x_g_* is the number of shared ancestors *g* generations ago. The vector {*x*} is, in a sense, a lower dimensional representation of the shared pedigree, and can be used to compute the probability of coalescence at each generation. For example, if *x*_1_ = 2 (full siblings), then all loci have the same 25% probability of coalescing within a single generation. We only consider the first *G* = log_2_ *N_e_* generations, where *N_e_* is the (constant) effective population size, as it was shown that the effect of the shared pedigree is important only up to ≈ log_2_ *N_e_* generations (Wakeley et al., 2012; Derrida et al., 2000; Chang, 1999). Beyond that time, almost all ancestors are shared, and the distribution of the contribution of each ancestor to the present day sample is approximately stationary.

By the law of total covariance, we have:

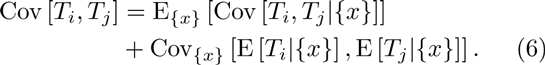

E_{*x*}_ [Cov [*T*_*i*_,*T*_*j*_|{*x*}]] ≈ 0, because conditioning on the pedigree, the loci are independently segregating. Therefore:

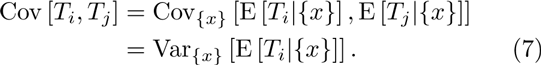

To compute E [*T*_i_|{*x*}], we condition on whether coalescence has occurred in the first *G* generations. If it has not occurred, we assume that the process then behaves just as the standard coalescent, or E [*T*_*i*_|no coal] = 2*N_e_* + *G*. We can write:

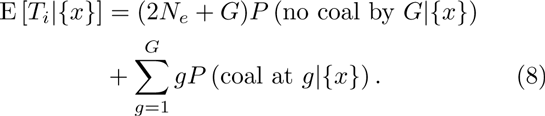

As computed in Wakeley et al. (2012), the coalescence probability is roughly given by 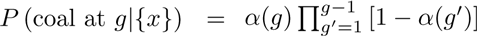, where *α*(*g*) = *x_g_*/2^2*g*+1^ and 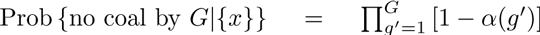. Since *α*(*g*) ≪ 1 (see below), we approximate *P* (coal at *g*|{*x*}) ≈ *α*(*g*) and 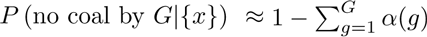. Thus,

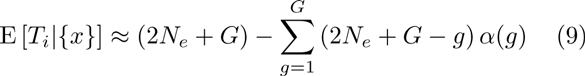

and

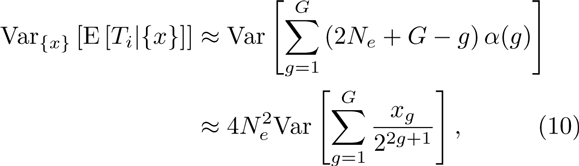

since *G* ≪ *N_e_*.

In Supplementary Material Section S1, we provide a numerical method to calculate the exact covariances of the *x_g_*’s under a diploid, discrete-time Wright-Fisher model (see the next section for definitions). To proceed here, we assume that the *x_g_*’s are independent. While the *x_g_*’s are clearly positively correlated, the independence assumption allows us to derive a lower bound on Cov [*T_i_*,*T_j_*], and thereby, the variance of 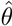. Under that assumption, Eq. (10) becomes

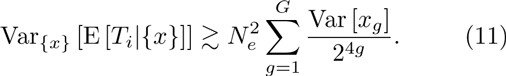

To compute the variance of *x_g_*, we note that the distribution of *x_g_* is roughly hypergeometric with parameters 2^*g*^ potential successes (the number of ancestors of one individual), *N_e_* − 2^*g*^ potential failures (all individuals in the population who are not ancestors of that individual), and 2^*g*^ draws (the number of ancestors of the other individual), giving 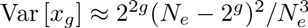. We provide the exact distribution of the variance of *x_g_* in Supplementary Material Section S1. Substituting the hypergeometric variance in Eq. (11),

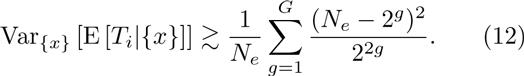

Using *G* = log_2_ *N_e_*, we have 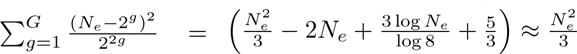 for large *N_e_*, and hence, using Eq. (7),

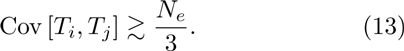

Using Eq. (4) and *θ* = 4μ*N_e_*, we finally obtain

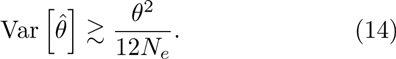

In summary, the variance due to the shared pedigree is of order *θ*^2^/*N*_e_, independently of the number of regions *n*. Thus, as argued above, even for a large number of chromosomes, the variance of 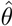 does not decay to zero, but rather to a constant that depends on the effective population size.

To intuitively explain the non-zero variance, we note that the pedigree itself is the product of a stochastic model (Wright-Fisher or another). Thus, even a fully specified pedigree, as obtained by sampling infinitely many loci, leaves uncertainty regarding the value of *θ*. In other words, the uncertainty in the estimate of *θ* results from having at hand only a single instance of a pedigree generated from the stochastic model governed by that parameter (see also Ralph (2015)).

## 4 Exact results for the correlation of the coalescence times at unlinked loci

In this section, we provide an exact derivation of the correlation of coalescence times at unlinked loci under a diploid, discrete-time, Wright-Fisher model. Further, we consider multiple sampling configurations for those loci, as explained below.

### 4.1 The sampling configurations

To compute the correlation of coalescence times at a pair of unlinked loci, we first note that there are multiple ways by which two such loci can be sampled in two individuals (or sequences). The six *sampling configurations* are shown in Figure 1. Four of these configurations involve a sample of two individuals, and we start by describing these.

**Figure 1:**
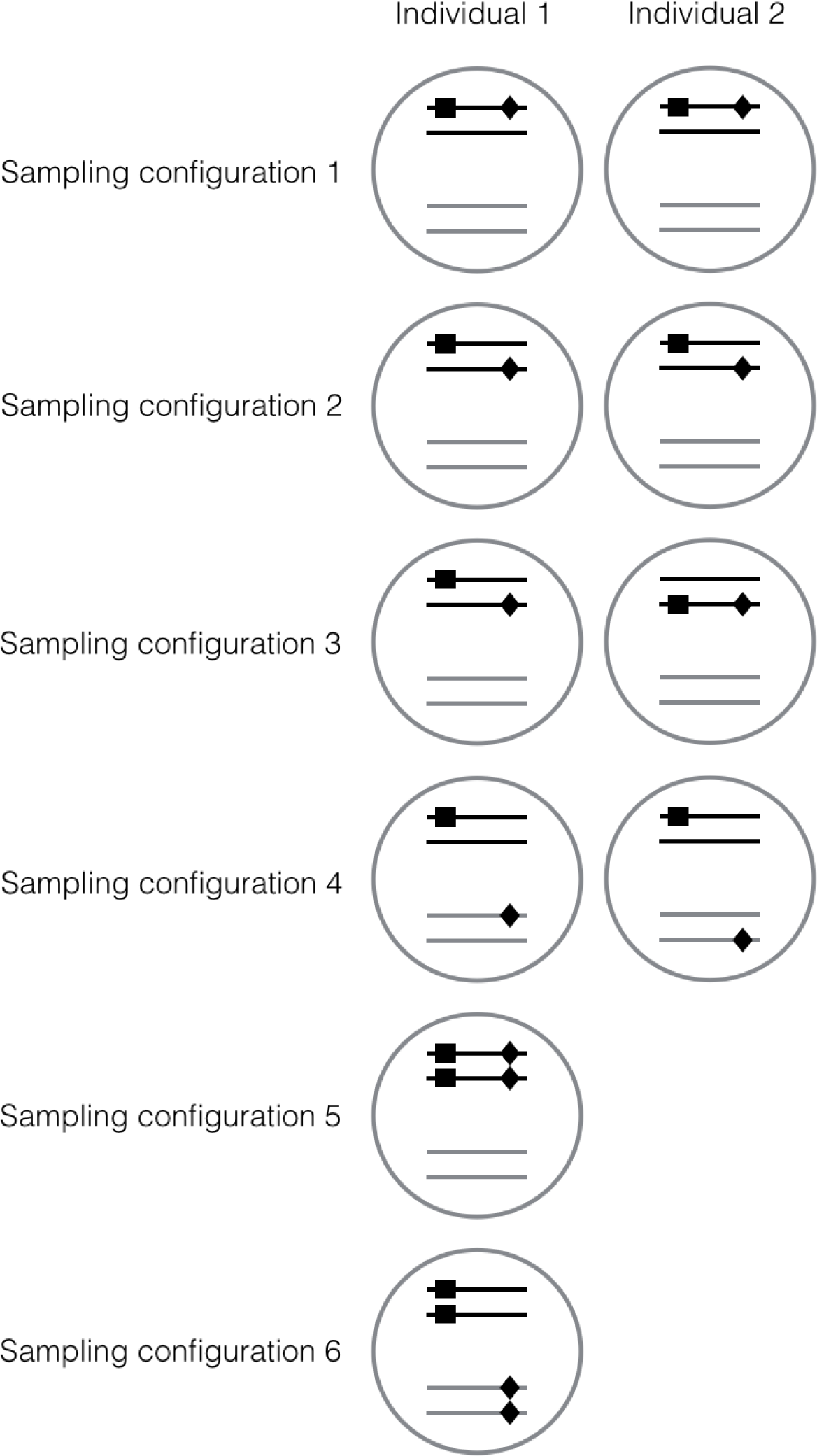
The sampling configurations. Sampling configurations 1 to 4 involve a sample of two individuals, depicted by two circles. Sampling configurations 5 and 6 involve a single individual, depicted by a single circle. The lines within each circle correspond to two pairs of homologous chromosomes. The two loci are indicated by squares and diamonds.

In the first configuration, the loci are located effectively infinitely far apart on the same chromosome in both individuals. This means that these loci will be coupled for the first few generations, until separated by a recombination event. Once separated, they may later back-coalesce onto the same chromosome, and again resume percolating together through the pedigree for a period of time that is expected to be short. (In the event of back-coalescence, two ancestral loci not sharing genetic material come to be located on the same chromosome, which essentially undoes the effect of recombination.) In the second configuration, the loci are on different homologous chromosomes, meaning they will necessarily be present in different parents in the immediately preceding generation, as each chromosome was inherited from a different parent. It is then also possible for them to back-coalesce in later generations. The third configuration is a mixture of the first two: the loci are located on the same chromosome in one individual, and on homologous chromosomes in the other. In the fourth configuration, the loci are sampled from non-homologous chromosomes in both individuals. This configuration is different from the previous three in that back-coalescence is not possible.

In the fifth and sixth sampling configurations, all sequences are sampled from a single individual. This is common in practice, as measuring the heterozygosity in a single individual does not require haplotype phasing. In configuration 5, we sample two loci from the same chromosome (and their pairs from the homologous chromosome). Given that each homologous chromosome must originate from a different parent, in one generation the sampled loci will transition to configuration 1 with probability 0.25, to configuration 2 with probability 0.25, and to sampling configuration 3 with probability 0.5. In sampling configuration 6, the sampled loci are on different (non-homologous) chromosomes. This configuration is reduced in one generation to sampling configuration 4, and therefore has the same correlation properties as that configuration.

### 4.2 The DDTWF model

To study the correlation of coalescence times under the different sampling configurations, we use a discrete-time Wright-Fisher (DTWF) model. This class of models has been advocated as an alternative to the coalescent when the sample size is large relative to the population size, as it can accommodate multiple and simultaneous mergers (Bhaskar et al., 2014).

In our case, we assume non-overlapping generations, a constant population size of *N_e_ diploid* individuals, half of which are males and half of which are females, random mating between the sexes, no selection, and no migration. There are three possible events: recombination, coalescence, and back-coalescence. Because the population size is finite, combinations of these events can occur in a single generation. We also keep track of whether lineages are in the same individual or not, as this determines their trajectory in the immediately preceding generation. We refer to this model as the 2-sex DDTWF. (Later, we also consider a simplified (1-sex) DDTWF). The dynamics of this 2-sex DDTWF model can be summarized by a Markov transition matrix (Supplementary Material Section S2) with 17 states, where the initial state is one of the sampling configurations 1, 2, 3, or 5.

The model described above represents pairs of loci sampled from either the same chromosome or homologous chromosomes, as the notion of back-coalescence and recombination only applies for these cases. Nevertheless, we found that the same transition matrix applies to sampling configurations 4 and 6 (non-homologous chromosomes), albeit with a different interpretation of the states (not shown).

Given the transition matrix, we can write a system of equations using a first step analysis for all states *x* such that *E*[*T_i_ T_j_*|*x*] > 0:

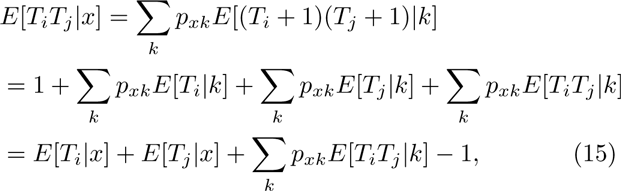

where *p_xk_* is the transition probability between states *x* and *k*.

Solving this system of equations allows us to obtain exact results for Cov [*T_i_*,*T_j_*|*x*]. As a note, *E*[*T_i_*|*x*] can be different from *E*[*T_j_*|*x*] depending on the state *x*. For example, if the pair of lineages at locus *i* is located on two different chromosomes in the same individual, whereas the pair of lineages at locus *j* is located in two different individuals, then *E*[*T_i_*|*x*] = *E*[*T_j_* |*x*] + 1. See more details in Supplementary Material Section S2. To obtain the correlation coefficient, we then normalize the covariance by the variance of the coalescence time at a locus, which is the same regardless of whether the lineages were sampled from the same or from different individuals. The variance can be calculated using the aforementioned system of equations with *i* = *j*.

Figure 2 shows the correlation coefficient of the coalescence times for each sampling configuration. The highest correlation is found for configuration 1. As the two loci are located on the same chromosome in both sampled individuals, they must have originated from the same parent in the previous generation. Therefore, both loci either both coalesce to the same parent or both do not, introducing correlation between the coalescence times. The effect of this sampling configuration then persists, as long as there is no recombination. As *N*_e_ increases, the correlation decreases, as it is much more likely for a recombination event to occur before a coalescence event. Sampling configuration 3 (two loci located far apart on the same chromosome in one individual, and on different chromosomes in the second individual) shows the lowest correlation. In fact, it is slightly negative for very small values of *N*_e_, for if one of the loci coalesces in the first generation, then it is impossible for the other locus to coalesce. The correlation in other configurations is intermediate between those of configurations 1 and 3.

**Figure 2:**
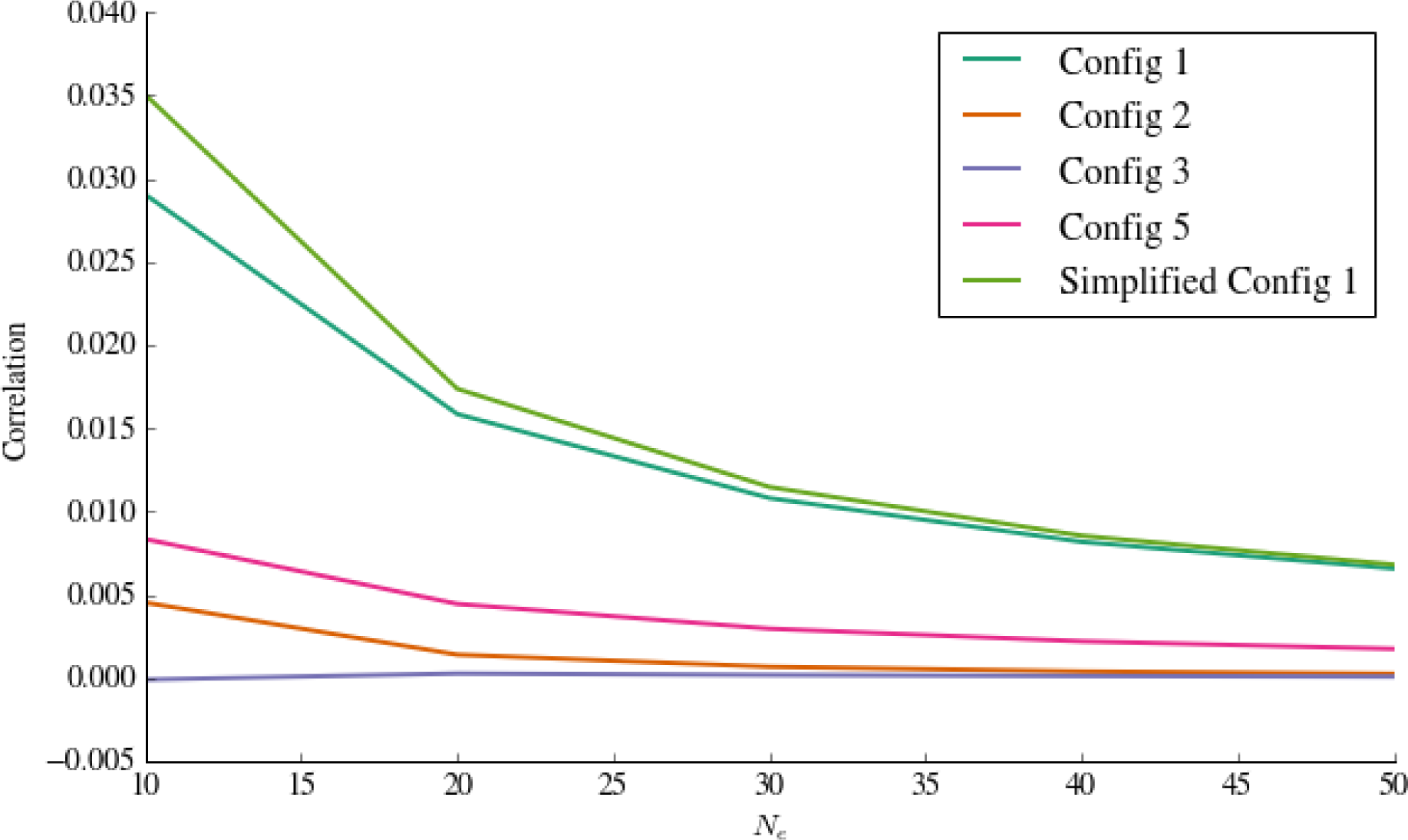
Correlation of coalescence times for a sample of size 2. We plot the correlation coefficients for the different sampling configurations under the 2-sex DDTWF and the simplified DDTWF vs the effective population size *N_e_*. The calculations are described in detail in Supplementary Section S2.

Figure 2 also shows results for a simplified DDTWF model, which is similar to the 2-sex DDTWF, except that individuals are monoecious and we do not keep track of whether lineages are in the same individual or not. There are fewer states in this model than in the 2-sex DDTWF, and it is therefore significantly easier to analyze. The simplified model displays a slightly higher correlation compared to the 2-sex model for *N_e_* ≲ 40, but is a good approximation otherwise (as we also show in Section 6). More details on both models are given in Supplementary Material Section S2.

## 5 Simulations

### 5.1 Wright-Fisher simulations

In this section, we use simulation of the 2-sex diploid, discrete-time Wright-Fisher model to support our analytical results from Section 3.2. To estimate the correlation coefficient of the coalescence times at two loci, we first simulate many Wright-Fisher pedigrees and sample, for each pedigree, two individuals from the current generation. We set the population size *N_e_* to be the same in every generation, with equal numbers of males and females. We then consider two loci on non-homologous chromosomes and simulate the path through the pedigree that connects the two lineages at each locus to their most recent common ancestor. In each generation and for each locus, lineages that are found in the same individual coalesce with probability 1/2, in which case the coalescence time is recorded. Loci on different chromosomes in the same individual coalesce neither in that generation nor in the previous generation.

We repeat this process multiple times for each pedigree to obtain an estimate of E[*T*|ped]. We then compute its variance over many simulated pedigrees to obtain Var_ped_ [E[*T*|ped]. By the same logic as Eq. (7) above, Var_ped_[E[*T*|ped]] is equal to Cov [*T_i_*,*T_j_*]. To obtain the correlation coefficient, we divide Cov [*T_i_*, *T_j_*] by Var [*T*] = Var_ped_ [E [*T*|ped]]+E_ped_ [Var [*T*|ped]]. The simulation results are shown in Figure 3. Our analytical lower bound, which, based on Eqs. (14) and (5), can be written as Corr [*T_i_*,*T_j_*] ≳ 1/(12*N*), is well supported by the simulations, and is in fact relatively tight.

**Figure 3:**
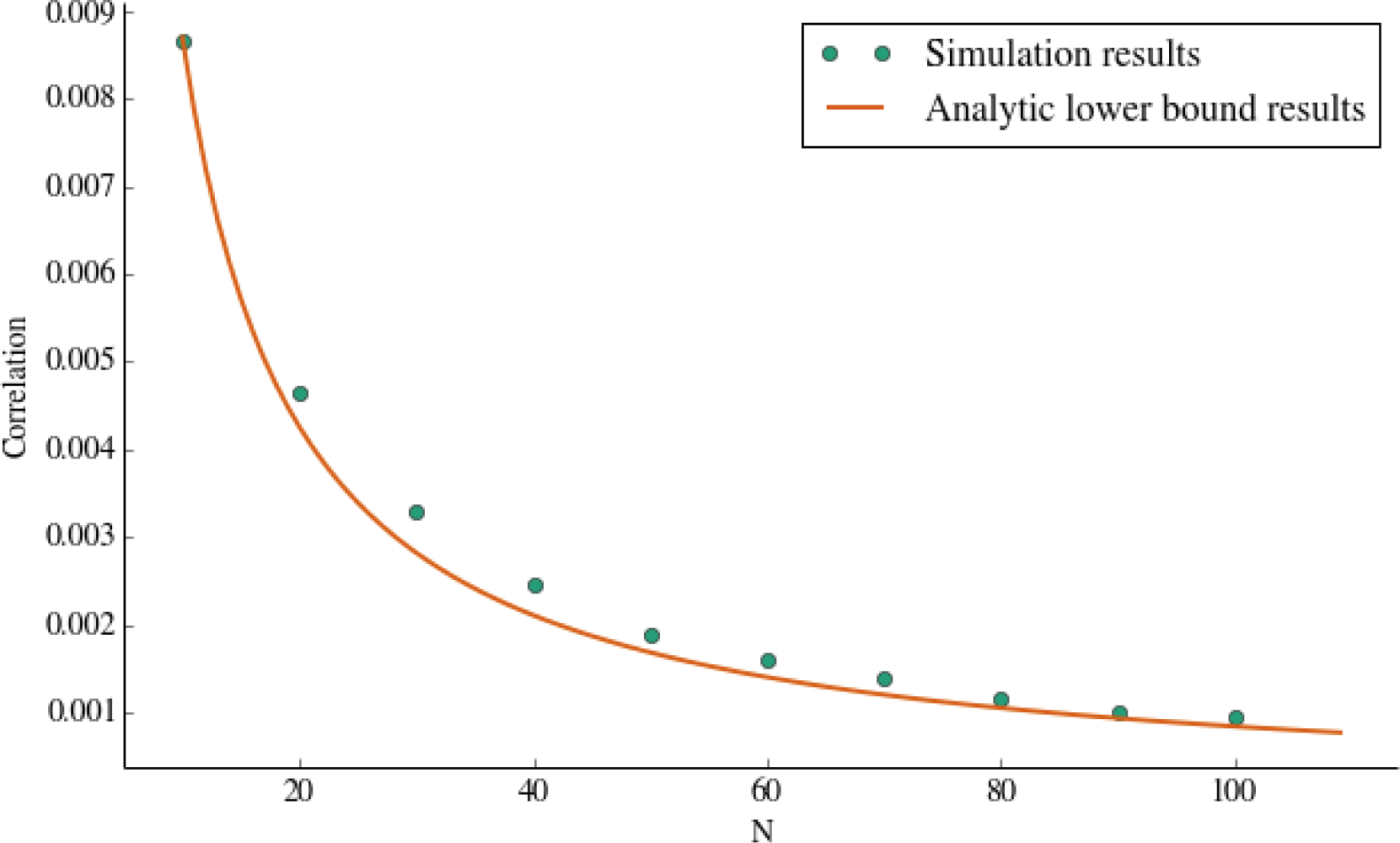
Analytical lower bound for the correlation of coalescence times at unlinked loci. We plot the correlation coefficient of the coalescence times at unlinked loci sampled from non-homologous chromosomes under the 2-sex, diploid, discrete-time Wright-Fisher model (green circles) as a function of the effective population size *N_e_*. The analytical lower bound (Corr [*T_j_*,*T_j_*] ≳ 1/(12*N*)) is plotted as a solid line.

### 5.2 Simulations based on real human pedigrees

The WrightFisher model is only one way to generate pedigrees having a given effective population size. In real human populations, pedigrees have complex structures that depend on their geographical region. For example, there are different rates of consanguineous marriages in different countries (Bittles and Black, 2015), different distributions of the number of children per family, and different mating structures, leading to differences in the number of full-siblings and half-siblings. To gain insight on the effect of these differences on the ability to estimate *θ*, we constructed a Wright-Fisher-like model, but which is constrained by patterns of real human pedigrees. Specifically, we used the Familinx database, compiled by Erlich (2016), which carries information on about 44 million individuals from different countries.

We extracted genealogical data for three countries (Kenya, Sweden, USA) from Familinx; these countries were arbitrarily selected among those with sufficient data. We then used these genealogies to simulate pedigrees by breaking down and reassembling small family units, as previously described for a different dataset (Wakeley et al., 2012). Specifically, we first split the genealogies into two-generational family units of children and their parents. To belong to a unit, a child must share at least one parent with at least one other child in the family unit. Because Familinx contains data on more than the three countries we chose, then in order not to create a bias in favor of smaller, simpler family units, we only require that the first sampled child be in the corresponding country data set. These family units then serve as building blocks to generate pedigrees with the same mating patterns and distribution of the number of children as in the reference population. Under these models, the effective population size *N_e_* is not guaranteed to equal the census population size. Therefore, to determine the effective population size for each model, we estimated *N_e_* as half the empirical average time until coalescence across randomly sampled pairs and random pedigrees. We could then fine-tune the census size, for each country, until reaching a pre-specified *N_e_*. Once the pedigrees were generated, we simulated genealogies through those pedigrees as described in Section 5.1. Additional details are provided in Supplementary Material Section S3.

For each country and for a range of *N_e_*’s, we then used the simulated data to compute the correlation coefficient of the coalescence times, as in Section 5.1 (i.e., Var_ped_ [E [*T*|ped]] divided by Var[*T*]). The results, shown in Figure 4, demonstrate that Corr [*T_i_*,*T_j_*], and consequently, 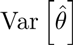, vary across populations, and are higher in the Famlllnx-inspired models compared to the Wright-Fisher model. A plausible explanation is that in the Wright-Fisher model, the ratio of half siblings to full-siblings is much higher than in the human pedigrees; this implies higher variance in the degree of relatedness in many real-world pedigrees relative to Wright-Fisher pedigrees. Therefore, it would be more difficult to estimate *θ* (i.e., the variance of 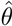 will be higher) in real-world populations than based on the expectation from the Wright-Fisher model. Further deviations are expected if we were to impose realistic first-cousin mating rates (Bittles and Black, 2015).

**Figure 4:**
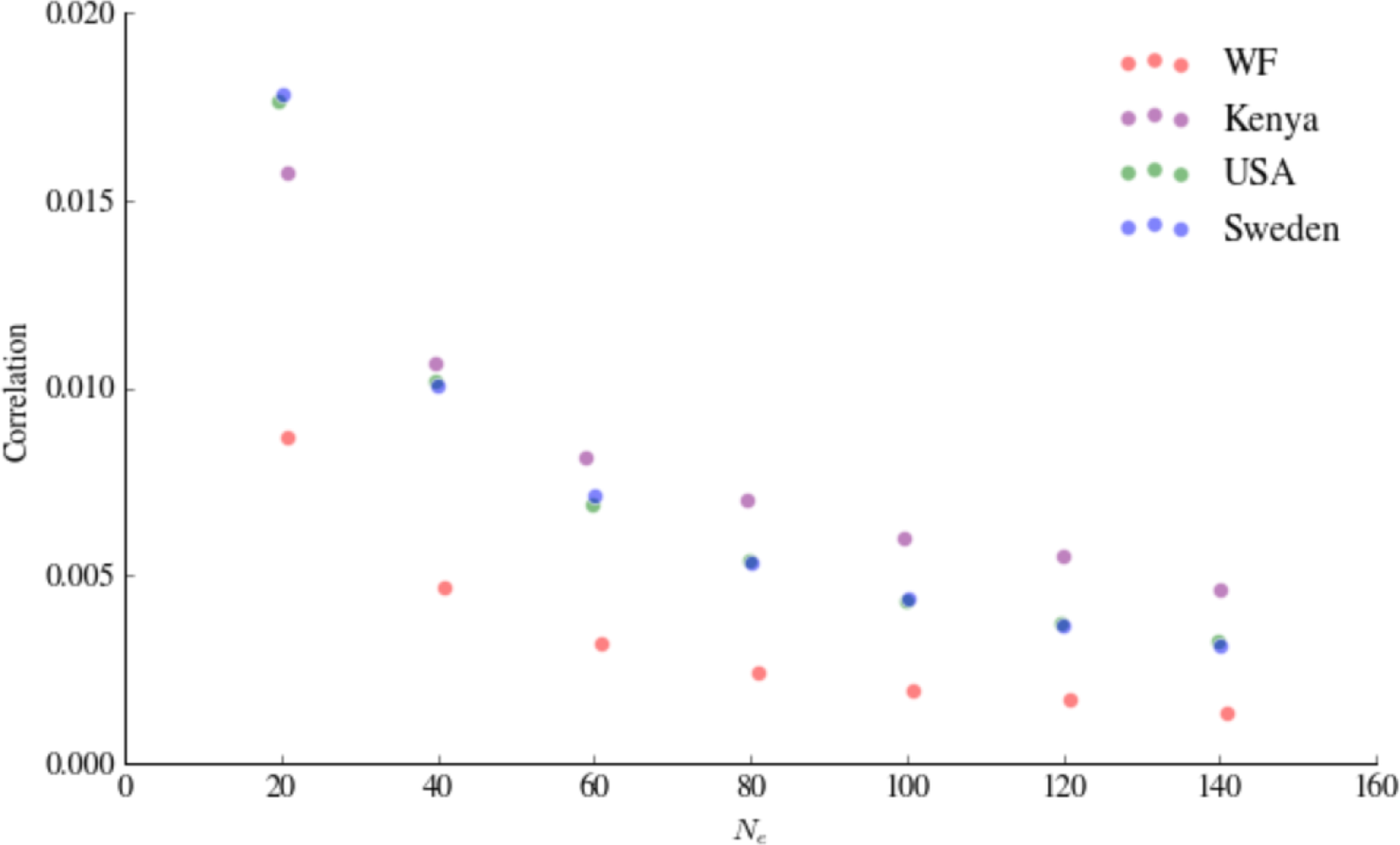
The correlation coefficient of coalescence times at two unlinked loci under synthetic pedigrees constructed using the Familinx dataset. Results are shown for three countries, as well as for the 2-sex DDTWF model. The correlation coefficient is plotted vs the effective population size *N_e_* (see the main text on how *N_e_* was set for the Familinx pedigrees). The two loci were sampled from non-homologous chromosomes. It can be seen that the correlation depends on the structure of the pedigree in ways that cannot be summarized by *N_e_*.

## 6 Linked sites and model comparisons

We have so far only studied unlinked sites; however, our analytical results for the DDTWF models can be relatively easily extended to the case of linked loci. Such an extension is important, since, for example, the covariance of coalescence times at two loci is directly related to the r^2^ measure of linkage disequilibrium (McVean, 2002). Quantifying the behavior of different models in terms of the covariance of coalescence times can thus provide insight into the importance of certain modeling assumptions.

In the DDTWF model with linked sites, the transition probabilities are expressed in terms of the per generation recombination probability, *r*, which has been so far set to 0.5. The transition matrix of Supplementary Material Section S2 is straightforward to adapt for any *r* < 0.5, and the covariance or correlation coefficient of the coalescence times can be computed. The correlation coefficient under the 2sex DDTWF model is plotted in Figure 5 vs the scaled recombination rate *ρ* = 4*N_e_r*, showing perfect agreement with simulations.

**Figure 5:**
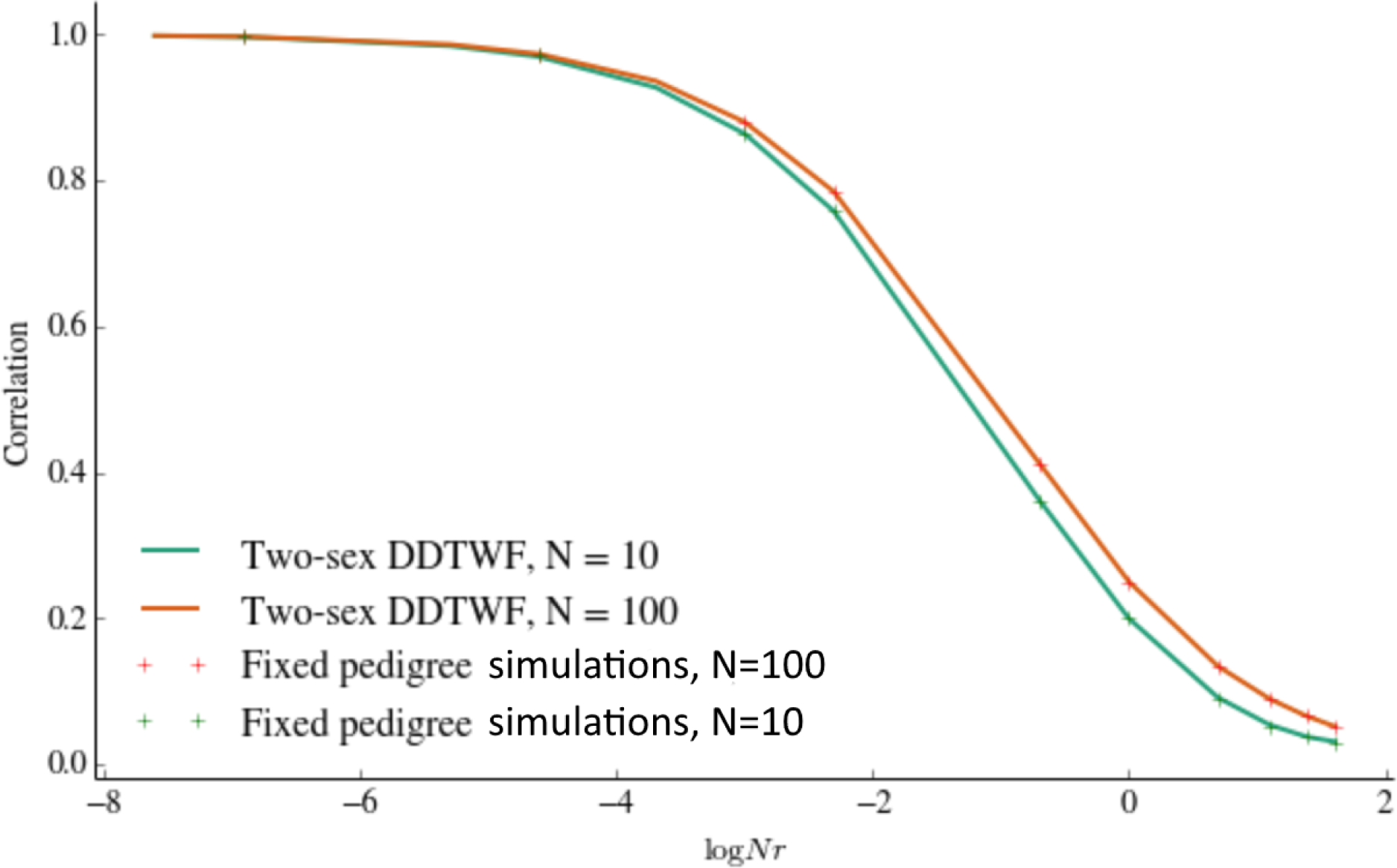
The correlation coefficient of coalescence times at two linked loci under the 2-sex DDTWF model. The correlation coefficients are plotted as lines for two values of *N_e_* vs the scaled recombination rate *ρ* = 4*Nr*. Simulation results are shown as + symbols. The two loci were sampled in configuration 1.

These results now enable us to compare the exact 2-sex DDTWF model to the simplifed DDTWF model, as well as to the coalescent with recombination and its Markovian approximations. Let *ρ* = 4*Nr*. Under the ancestral recombination graph (ARG) (Griffiths and Marjoram, 1997), which is the standard model for the coalescent with recombination, the covariance of coalescence times at two loci satisfies (e.g., Simonsen and Churchill (1997)),

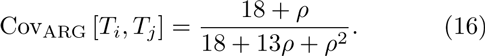

Under the Sequentially Markov Coalescent (SMC) (McVean and Cardin, 2005), each new genealogy (following recombination) depends only on the previous genealogy (as opposed to the ARG (Wiuf and Hein, 1999)), and the new coalescence time must differ from the previous time (no back-coalescence allowed). In this case, we have,

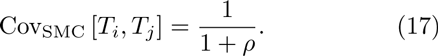

The SMC’ model (Marjoram and Wall, 2006) is a variant of SMC where back-coalescence is allowed. Under SMC’ (Eriksson et al., 2009; Wilton et al., 2015),

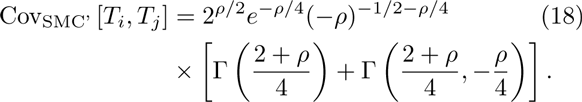

(The covariances of Eq. (16)-(18) are also equal to their respective correlation coefficients, since Var [*T*] = 1 under either the ARG, SMC, and SMC’). In Figure 6, we compare the correlation of *T_i_* and *T_j_* across the different models as a function of *ρ* for *N_e_* = 100 and different values of *r*. The ARG provides a very good approximation under these conditions. In turn, the SMC’ model shows very slight deviations compared to the ARG, while, as previously shown, the SMC model deviates more substantially (Wilton et al., 2015).

**Figure 6:**
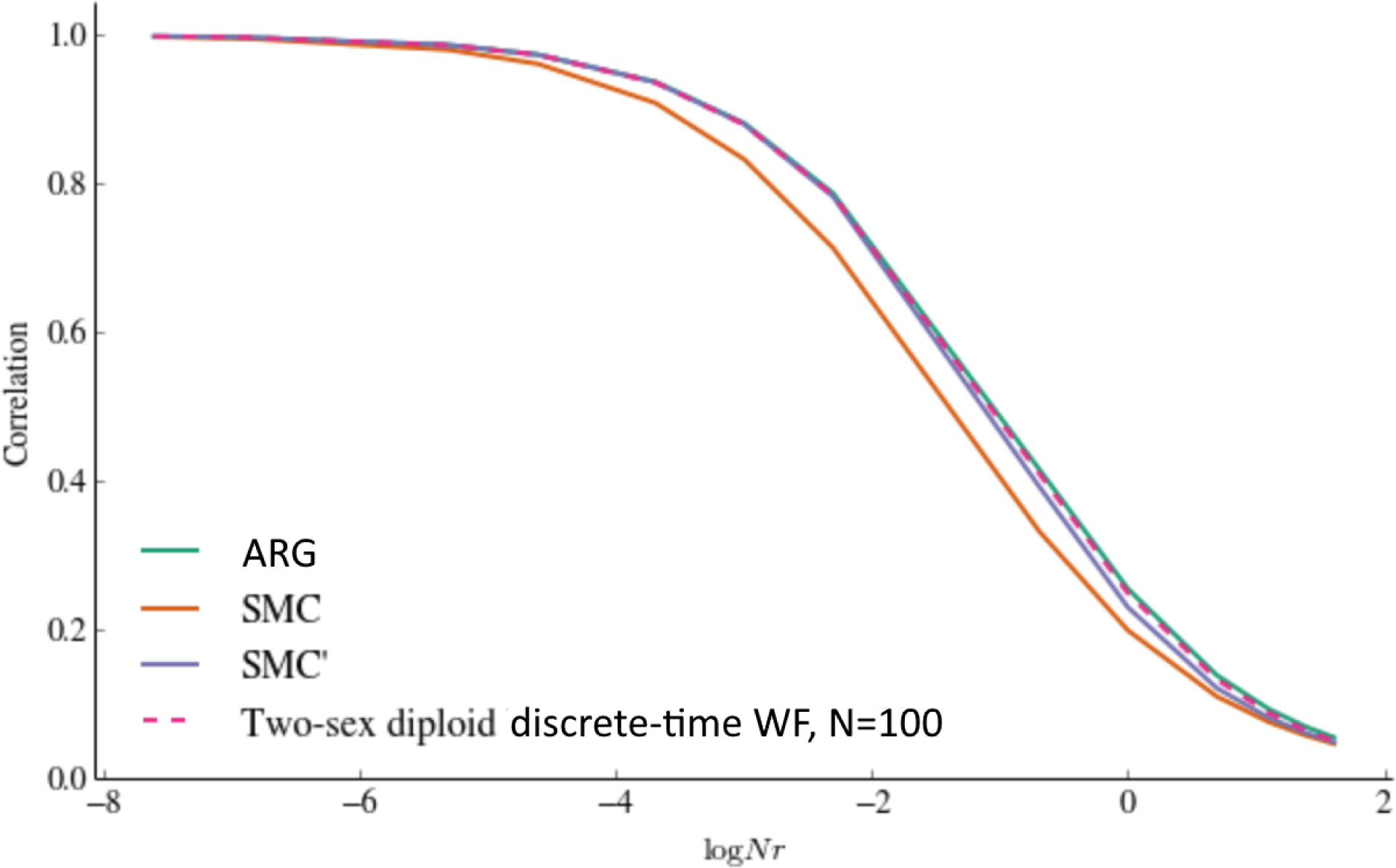
A comparison of the correlation coefficient of the coalescence times at two linked loci under models of increasing complexity. We compare the ARG, SMC, SMC’, and the 2-sex DDTWF with *N_e_* = 100, across different values of *ρ* = 4*N_e_r*. The predictions of the ARG and SMC’ are very good approximations for those of the 2-sex diploid Markov Chain model (for the value of *N_e_* shown here).

The 2-sex DDTWF model is compared to the simplified DDTWF model in Figure 7. Compared to the full 2-sex model, the simplified model is an extremely good approximation even for *N_e_* as small as 100: the maximum difference in the correlation coefficient (across different values of *r*) between these two models was less than .005 (see also Figure 2). Therefore, the simplified model should be preferred due its much reduced complexity. For *N_e_* = 10, we observe a more noticeable difference between the 2-sex and the simplified DDTWF models, with a maximal difference around .025.

**Figure 7:**
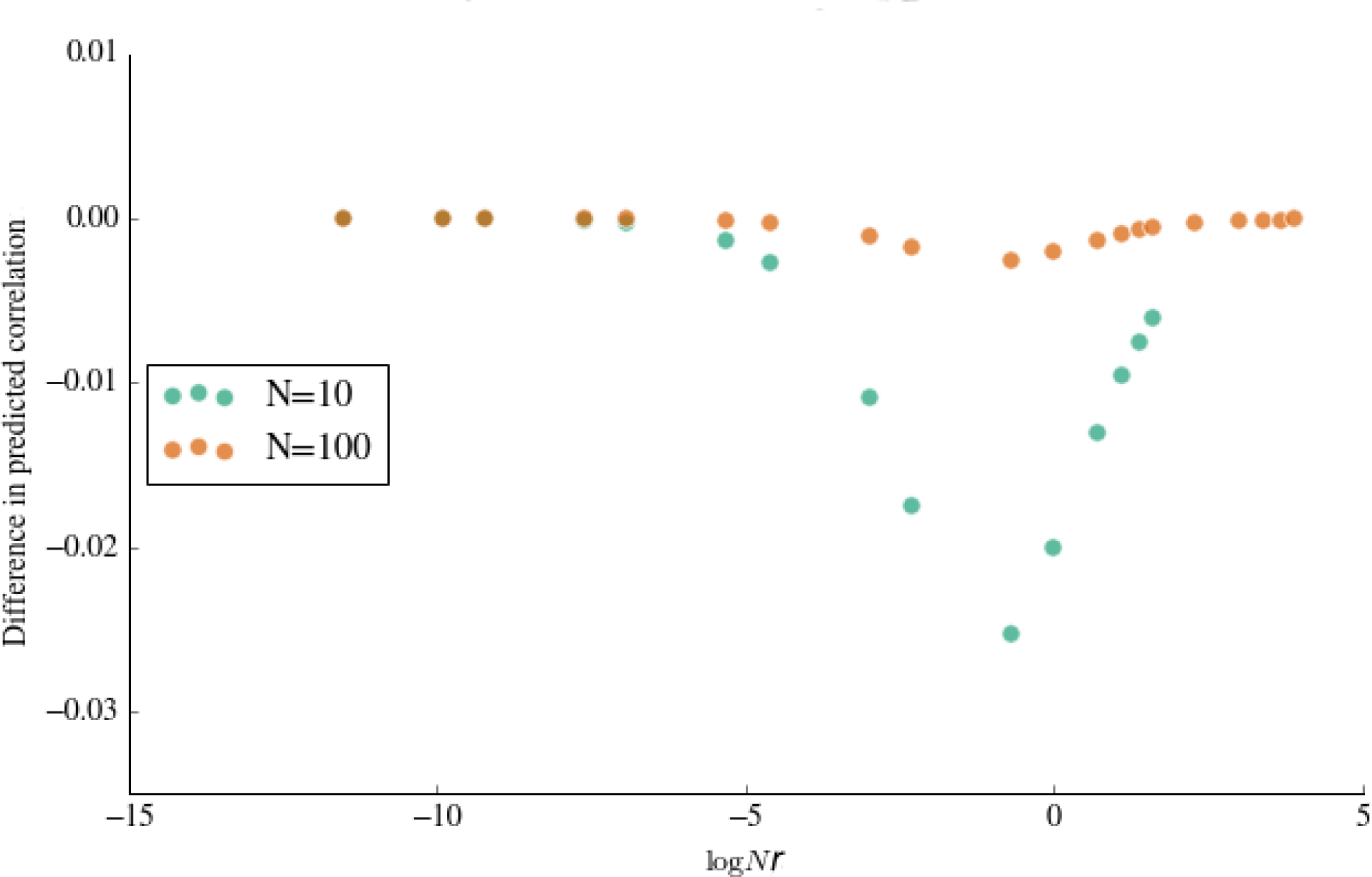
A comparison of the correlation coefficient of the coalescence times at two linked loci between the 2-sex and the simplified DDTWF models. We plot the difference between the correlation coefficients of the two models for *N_e_* = 10 and *N_e_* = 100 and for different values of *r*. The predictions of the two models slightly diverge at *N_e_* = 10.

## 7 Discussion

Increasing the size of the sample is known to have limited ability to improve estimates of *θ*, as the individuals in the sample share most of their genealogy (Rosenberg and Nordborg, 2002). For this reason, it was recommended to use data from many unlinked gene loci from a small number of individuals (Felsenstein, 2006). While this intuition still holds, we have shown that the estimator of *θ* based on the average number of pairwise differences at many loci is not consistent and has non-zero variance, even when sampling infinitely many loci. We have provided an approximate lower bound for the variance for loci on non-homologous chromosomes, as well as exact results for diploid, discrete time Wright-Fisher models under various configurations of two sampled loci.

Fundamentally, the non-zero variance of 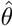 is a result the underlying pedigree shared between all loci. The shared pedigree itself is assumed to be a single draw from a random demographic process (Wright-Fisher or another), with a characteristic effective population size. Thus, even if we were able to perfectly characterize the single pedigree at hand, we cannot hope to infer with complete certainty the parameters of the demographic model. It is worth noting that one can adopt a different (philosophical) view, under which the pedigree itself is the subject of inference, and is not a product of a random demographic process (Ralph, 2015). Under such a view, there is no such thing as an estimator of the effective population size.

The analytical results in this paper are based on the Wright-Fisher model. To gain insight on the behavior of more realistic demographic models, we adapted the Wright-Fisher model according to the family structure of real human populations. The results demonstrated that the correlation of coalescence times is higher in the human-inspired models than in the WF model; therefore, *θ* should be more difficult to estimate than expected under the pure WF model.

When using a demographic model, it is not always clear which features of the real population are crucial (e.g., two sexes, diploidy, etc.), or whether simplified models could display similar characteristics. We used our analytical framework to study the correlation of coalescence times as a function of the scaled recombination rate, *ρ*, for the 2-sex and the simplified DDTWF models, and compared the results to the coalescent with recombination and its Markovian approximations. We found that, as expected, for sufficiently large effective population size (*N* ≳ 100), the results for the coalescent (as well as for its SMC’ approximation, but not for SMC) were extremely close to those of the DDTWF models. In contrast, differences were observed for *N* = 10, even between the 2-sex and the simplified DDTWF.

We have focused here on a sample of two individuals at two loci. For unlinked loci, we showed that the variance of 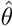 for any number of loci is reduced to the two-loci problem. Extending the sample size to more than two individuals is expected to be significantly more complicated. Deviations between the coalescent and the discrete time haploid Wright-Fisher model for increasing sample sizes were recently studied and shown to be important for realistic human demographic histories (Bhaskar et al., 2014). We similarly speculate the presence of a shared pedigree to have an increasingly significant effect on the variance of Tajima’s estimator as the sample size grows, but this analysis is left for future studies.

## Supplementary Material: Extended methods and analytical results

### S1 The number of shared ancestors

In this section, we derive the covariance of the number of shared ancestors at each generation for the 2-sex diploid discrete-time Wright-Fisher model (DDTWF), denoted *x_g_* in section 3.2 of the main text. The model is defined in Section 4.2 of the main text and Section S2 below. We proceed in three steps.

#### S1.1 Distribution of the number of ancestors from one generation to the next

Consider a single individual in a population with non-overlapping generations in the 2-sex model. Each generation *g*, there are *N_f_* males and *N_m_* females, where *N_f_* + *N_m_* = *N_e_*, and typically *N_f_* = *N_m_* = *N_e_*/2. Let *y_g_* be the number of ancestors of a particular individual at generation *g* in the past. During the first few generations, the number of ancestors grows very fast, and we expect *y_g_* ≈ 2^*g*^. As the number of ancestors in a given generation starts to approach the size of the population, the ancestors overlap with one another, and the growth of ancestors slows down until an equilibrium distribution is reached. We are interested in modeling the exact distribution of the number of ancestors in generation *g* + 1, *y_g_*+1, given the number of ancestors in generation *g*, *y_g_*.

We can first divide the number of ancestors in generation *g* + 1 into males and females:

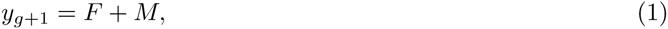

where *F* is the number of fathers of individuals in *y_g_*, and *M* is the number of mothers of individuals in *y_g_*. We have

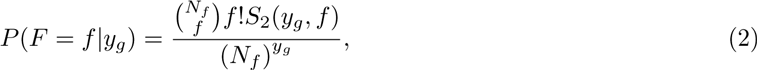

where *S_2_* is the Stirling number of the second kind. The intuition behind this formula is that there are 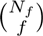 possible ways of choosing *f* fathers among the *N_f_* available. There are then *f*! possible orderings of these chosen males. The Stirling number of the second kind is the number of ways we can partition a set of *y_g_* individuals into *f* categories. We divide all this by the total number of ways of making *y_g_* choices of fathers among the *N_f_* available, or (*N_f_*)^*y_g_*^. Likewise,

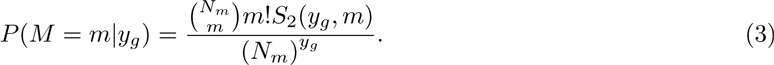

We then obtain the following convolution for the number of ancestors *a* in generation *g* +1,

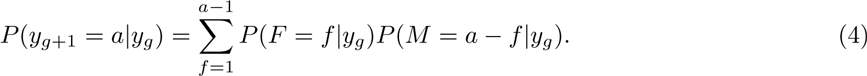

The numbers *y_1_*,…,*y_G_* form a Markov Chain. The preceding formula defines the transition matrix of *y_g_*+1 given *y_g_*.

If we did not have a 2-sex model, but instead a bi-parental monoecious model, the formula for the number of ancestors in generation *g* + 1 would be the following simpler expression

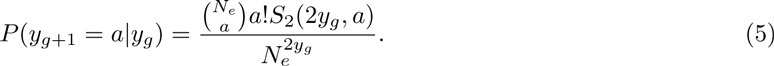

#### S1.2 Overlap in the number of ancestors each generation

In the previous subsection, we described the distribution of the number of ancestors at each generation. Here, we start with a sample of two individuals, A and B, and are interested in the distribution of the number of shared ancestors each generation. If this sample consists of a pair of full siblings, then the number of shared ancestors grows according to the formula provided in the previous section, as full siblings share all of their ancestors.

Let *X_g_* be the set of common ancestors in generation *g*, *A_g_* be the ancestors of *A* that are not in *X_g_*, and *B_g_* the set of ancestors of B that are not in *X_g_*. Let |*A*_*g*_|, |*B_g_*| and |*X_g_*| (= *x_g_* in the notation of the main text) be the cardinality of these three disjoint sets. Let *F_A_* be the set of fathers of individuals in *A_g_*, and let |*F_A_*| be the cardinality of *F_A_*. Likewise, we define *F_x_*, *F_B_*, |*F_X_*|, and |*F_B_*|. Given |*A_g_*|, |*B_g_*|, and |*X_g_*|, the distribution of |*F_A_*|, |*F_B_*|, and |*F_X_* | is as described in the previous subsection,

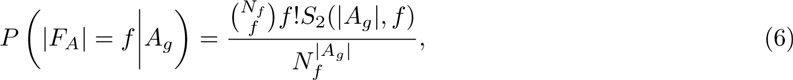

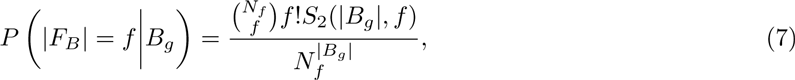

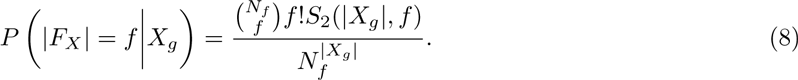

The number of fathers in common between individuals in *A_g_* and *X_g_*, *x_a_*, follows a hypergeometric distribution with *F_X_* success states, *N*_*f*(*g*+1)_ − |*F_X_*| failure states, and |*F_A_*| draws,

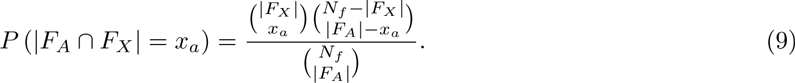

The probability that individuals in *B_g_* have *x_b_* fathers in common with individuals in *X_g_*, and *a_b_* fathers in common with individuals in *A_g_* (but not with individuals already in *X_g_*), given that |*F_A_* ∩ *F_x_* | = *x_a_*, is defined by a trivariate hypergeometric distribution,

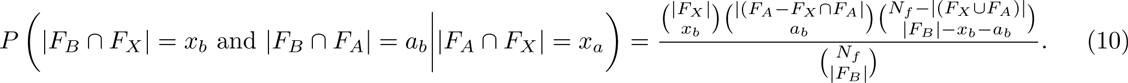

The number of shared male ancestors in generation *g* + 1 is |*X*_*f*(*g*+1)_| = |*F*_*X*_ | + *a_b_*, the number of male ancestors exclusive to A is |*A*_*f*(*g*+1)_| = |*F*_*A*_| − *a*_*b*_ − *x_a_*, and the number of male ancestors exclusive to B is |Bf(g+i)I = IFB I − ab − xb. To obtain the number of shared female ancestors, |*X*_*m*(*g*+1)_|, we use the same protocol, except replacing *N_f_* by *N_m_*. Finally, to derive the joint distribution of *X*_*g*+1_, *A*_*g*+1_ and *B*_*g*+1_, we take the convolution over the number of male and female ancestors.

In this way, we can derive a transition matrix *T*. The entries *T_ij_* of the transition matrix give the probability of entering state *j* = (|*A*_*g*+1_|, |*B*_*g*+1_|, |*X*_*g*+1_|) given state *i* = (|*A_g_*|, |*B_g_*|, |*X_g_*|).

We plot the dynamics of the number of shared ancestors along the generations in Supplementary Figure 1. The distribution of the number of shared ancestors in generation *g* is obtained by considering the *g*-th power of *T*, assuming a sampling configuration of (1,1, 0) and then summing over the probabilities of all configurations with same |*X_g_*|.

**Supplementary Figure 1:**
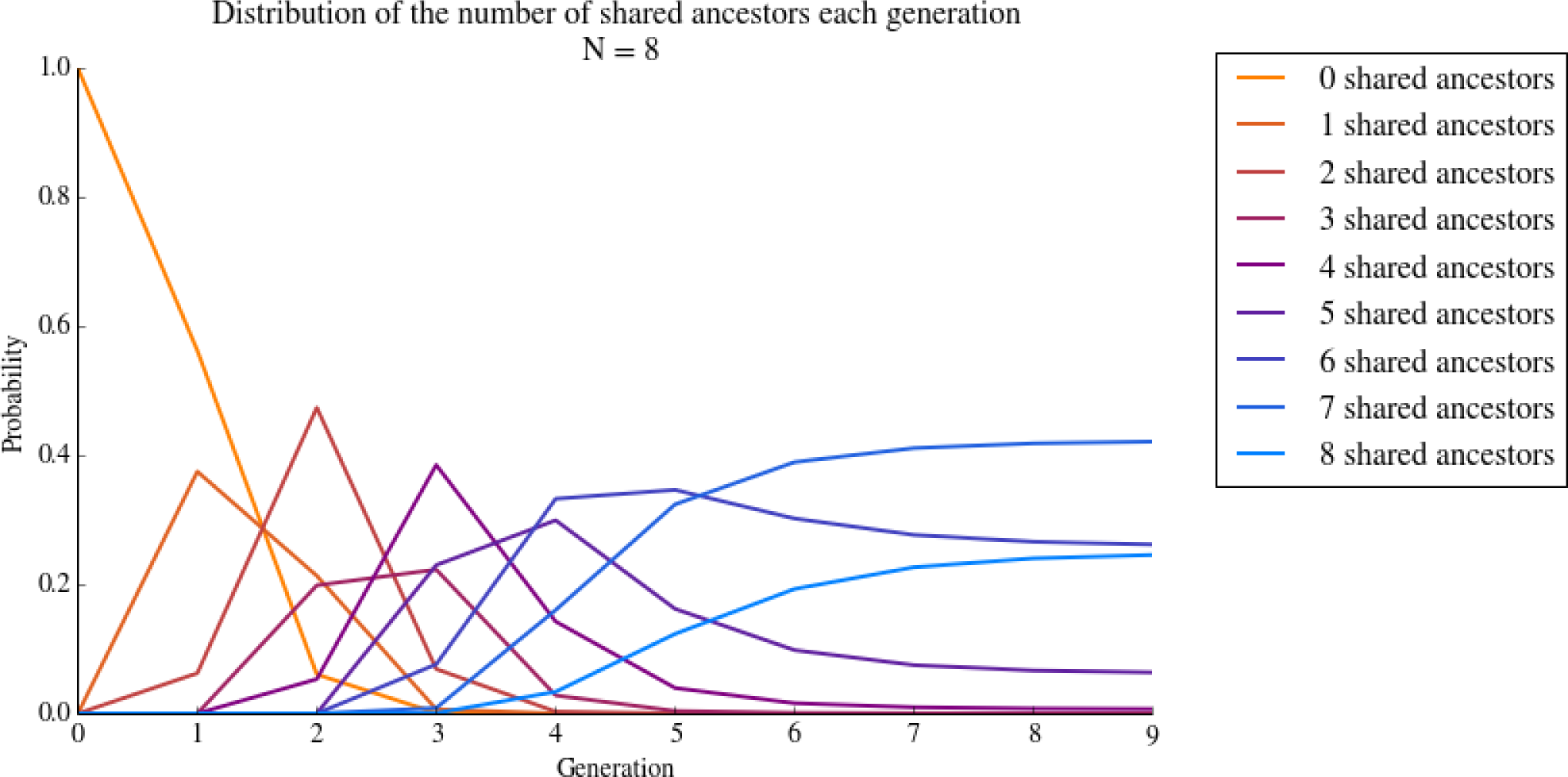
The distribution of the number of shared ancestors in each generation for the 2-sex DDTWF model. We used *N_e_* = 8. The process reaches an equilibrium distribution after about 7 generations.

Finally, consider the simpler bi-parental monoecious model, and let *K_A_*, *K_B_*, and *K_X_* be the parents of individuals in *A_g_*, *B_g_*, and *X_g_*, respectively. As in the previous subsection,

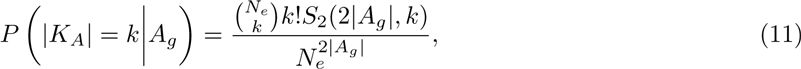

and similarly for *K_B_* and *K_X_*. As above, the number of parents in common between individuals in *A_g_* and *X_g_*, *x_a_*, follows a hypergeometric distribution,

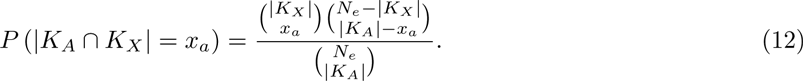

The number of parents common to *B_g_* and *A_g_* or *X_g_* is similarly given by

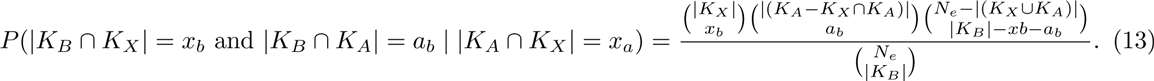

As above, we have |*X*_*g*+1_| = |*K_X_*| + *a_b_*, |*A*_*g*+1_| = |*K_A_*| − *a_b_* − *x_a_*, and |*B*_*g*+1_| = |K_B_| − *a_b_* − *x_b_*. This fully specifies the distribution of the configuration in generation *g* + 1, given the configuration in generation *g*.

#### S1.3 Variance and covariances of the number of ancestors each generation

Finally, we calculate the covariances between the number of shared ancestors in generations *i* and *j*, Cov(*x_i_*,*x_j_*), using the transition matrix *T* derived as described in the previous section. Let state 0 be the index of the sampling configuration, (1,1,0). We have, for *i* ≤ *j*,

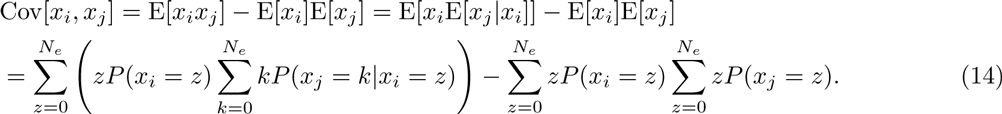

Each value of the number of shared ancestors, *z* is represented by multiple states of the transition matrix. We refer to the set of these states as “Conf *z*”, or

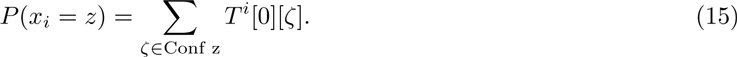

Thus,

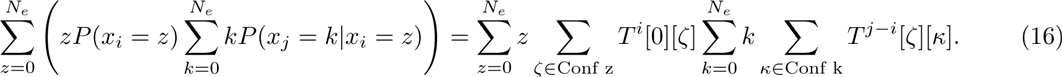

We plot the covariances and correlations for the 2-sex DDTWF model and for *N_*e*_* = 10 in Supplementary Figure 2.

**Supplementary Figure 2:**
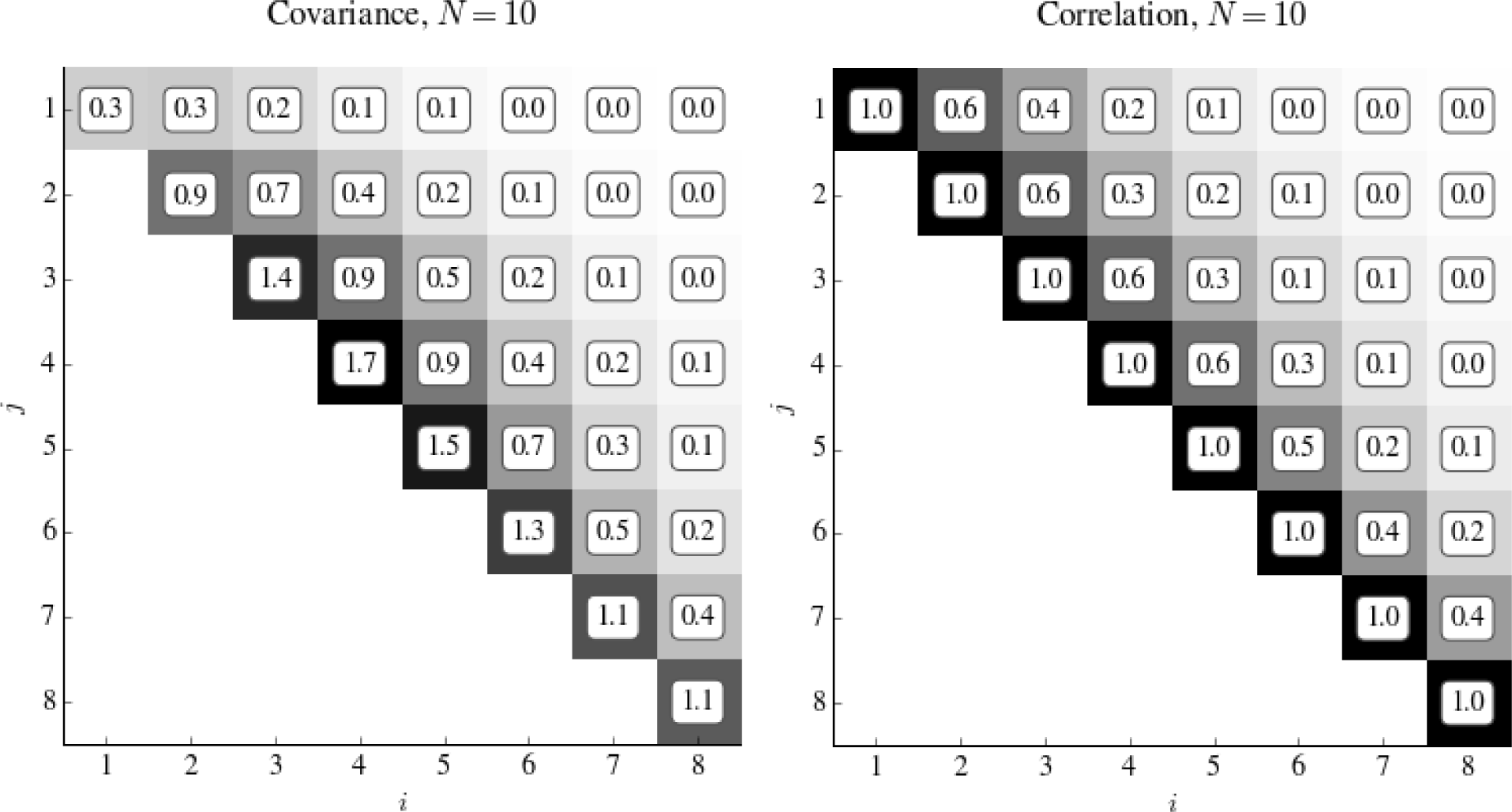
The covariance and correlation of the number of shared ancestors across the generations. In the left panel, we show the covariance of the number of shared ancestors, *x_g_*, for each generation *g*, and for *N_e_* = 10. The diagonal represents the variance of the number of shared ancestors, and is highest in generations 3-5. In the right panel, we show the correlation coefficients. The correlation between *x_g_* and *x*_*g*+1_ decreases with *g*.

We note that the entire derivation of this section can be generalized to the case when the number of males and females is allowed to differ as well as change along the generations.

### S2 The DDTWF models

#### S2.1 The 2-sex DDTWF model and transition matrix

The notation we use to label the states in this transition matrix is derived from the notation of Wakeley and Lessard (2003), who used a similar transition matrix to analyze patterns of linkage disequilibrium in a 2-locus multi-deme model. The notation is explained in Supplementary Figure 3. For example, state {12,12} represents the case where two copies of the first locus are located in two different individuals, and on the same chromosome as this first locus is the second locus. The comma separates the different chromosomes on which genetic material os tracked, and the numbers 1 and 2 represent the loci on each chromosome.

**Supplementary Figure 3:**
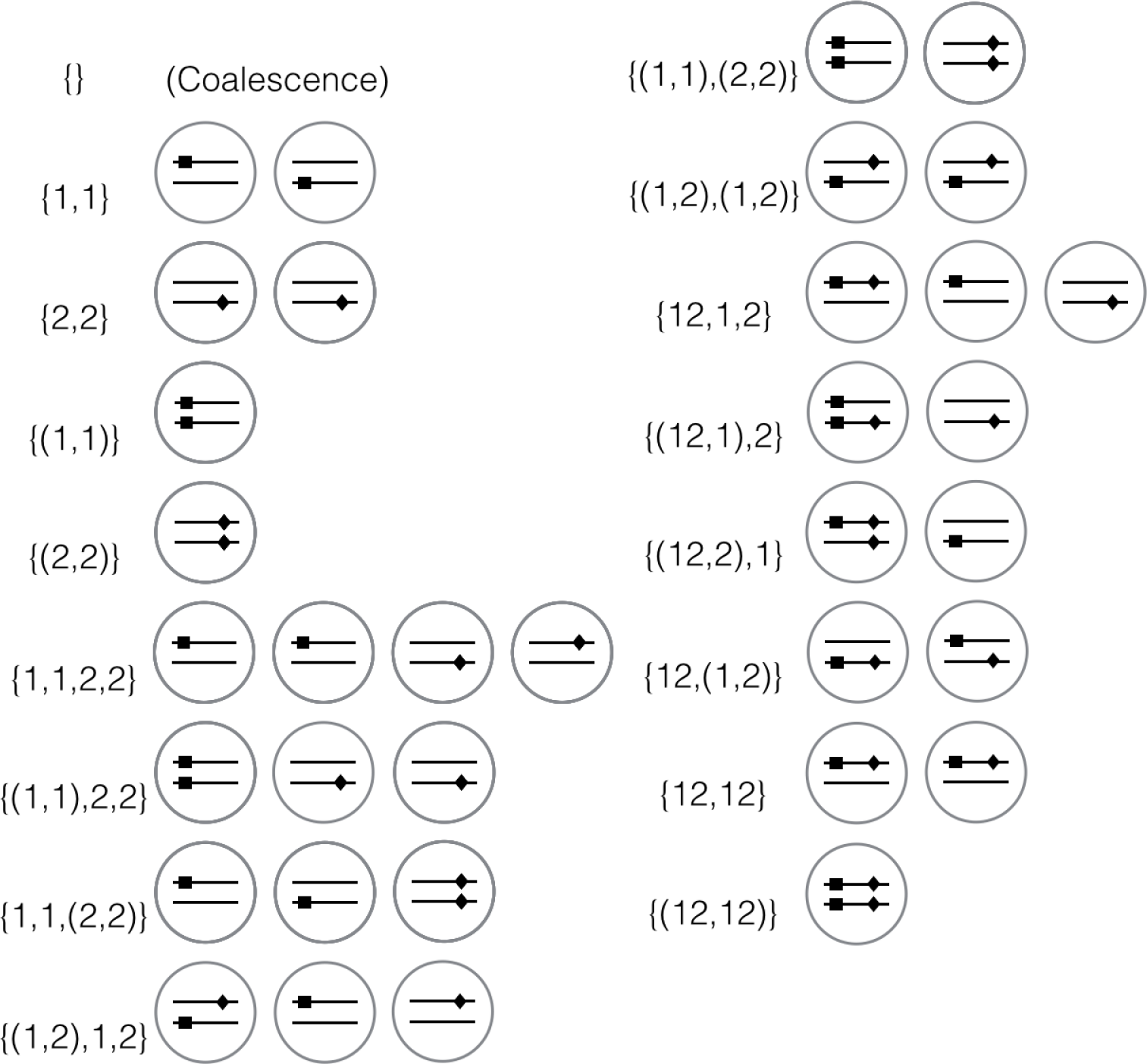
The states of the 2-sex DDTWF model. Circles represent individuals; the two lines within each individual represents a pair of homologous chromosomes; the square represent the first locus and the diamond represents the second locus. For example, {12,12} corresponds to the sampling configuration 1 in main text Figure 1.

In state {(12,12)}, the parentheses indicate that the tracked pairs of loci are present on two different chromosomes in the same individual. If the tracked lineages are on different chromosomes of the same individual, then they must be located in different individuals in the previous generation. So, for example, state {(1,1)} transitions to state {1,1} in one generation with probability 1.

The set of all possible states in our model is : {}, {1,1}, {2,2}, {(1,1)}, {(2,2)}, {1,1,2,2}, {(1,1),2,2}, {1,1,(2,2)}, {1,2,(1,2)}, {(1,1),(2,2)}, {(1,2),(1,2)}, {12,1,2}, {(12,1),2}, {(12,2),1}, {12,(1,2)}, {12,12} and {(12,12)}. We show the communicating states in this transition matrix in table S2.1.

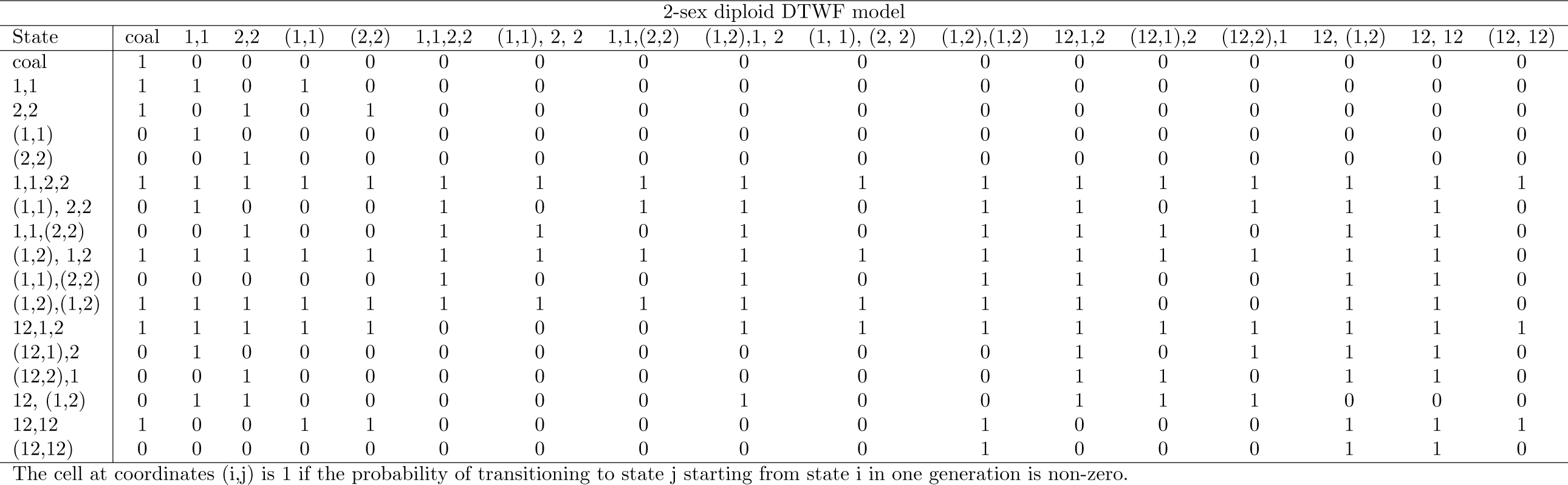

#### S2.2 The simplified DDTWF model

We also consider a simplified version of this model, a monoecious bi-parental DDTWF model. In this model, we do not keep track of whether lineages are in the same individual or not. The diploidy only comes into play in that recombination is impossible in a haploid context. The complete list of states in this model is: {}, {1,1}, {2,2}, {1,1,2,2}, {12,1,2}, and {12,12}, far fewer than in the 2-sex DDTWF model. For this reason, this model can only be used to show the effect of a limited number of sampling configurations. For example, it is not possible to model sampling configuration 2, where loci are sampled from different homologous chromosomes in the same individual. We show a matrix of communicating states in table S2.2.

| Simplified diploid DTWF model |  |  |  |  |  |  |
| --- | --- | --- | --- | --- | --- | --- |
| State | coal | 1,1 | 2,2 | 1,1,2,2 | 12,1,2 | 12, 12 |
| coal | 1 | 0 | 0 | 0 | 0 | 0 |
| 1,1 | 1 | 1 | 0 | 0 | 0 | 0 |
| 2,2 | 1 | 0 | 1 | 0 | 0 | 0 |
| 1,1,2,2 | 1 | 1 | 1 | 1 | 1 | 1 |
| 12,1,2 | 1 | 1 | 1 | 1 | 1 | 1 |
| 12,12 | 1 | 1 | 1 | 1 | 1 | 1 |

The cell at coordinates (i,j) is 1 if the probability of transitioning to state j starting from state i in one generation is non-zero.

#### S2.3 The expected generation time in both models

If two lineages are located in two different individuals, then the probability they coalesce in a single generation is just 1/(2*N_e_*). However, if they are present in different chromosomes of the same individual, they must have originated from two different individuals in the previous generation. Because of this, the expected time until coalescence will be different than 2*N_e_* in the 2-sex DDTWF, as opposed to in the simplified DDTWF where it is just equal to 2*N_e_*.

The process retains some memory of the fact that lineages were initially sampled in two different individuals. Indeed, the time until coalescence at generation *g*, given no coalescence in any previous generation, will be different than the expected time until coalescence at generation *g* + 1, given no coalescence in any previous generation. As *g* increases, this difference in coalescence times decreases from one generation to the next, and the process converges to an average generation time.

Consider a pair of lineages in two individuals. In generation *g* +1, given that no coalescence events have occurred in any of the previous *g* generations, the probability of the two lineages to coalesce is

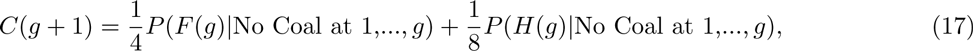

where *P*(*F*(*g*)|No Coal at 1,…,*g*) and *P*(*H*(*g*)|No Coal at 1,…,*g*) are the probabilities that the two lineages are located in full siblings and half siblings, respectively, in generation *g*, given no coalescence in that generation or any of the previous generations. Next, we write

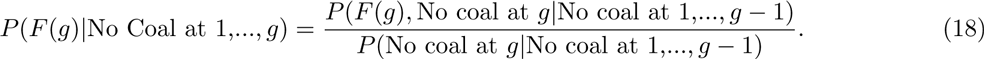

The denominator is simply given by 1 − *C*(*g*). For the numerator, we note that for the two lineages to arrive at full siblings in generation *g*, then first, we must exclude the possibility that the lineages are at the same individual in generation *g* (given no previous coalescence), which happens with probability 2*C*(*g*) (since the probability of coalescence is half the probability to arrive at the same individual). Second, the probability that these two individuals share both parents is 1/(*N_e_*/2)^2^. Therefore,

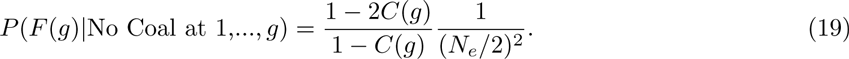

In the same way, we have

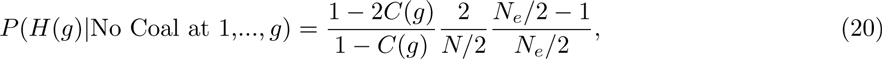

By solving *C*(*g* + 1) = *C*(*g*), we obtain the limiting coalescence probability as a function of *N_e_*. As the equilibrium distribution of the time until MRCA is geometric, the equilibrium generation time is the inverse of the probability of coalescence each generation, or

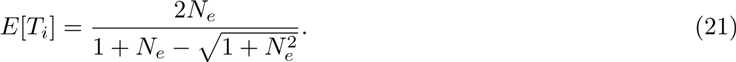

This generation time is always slightly greater than 2*N_e_*.*E*[*T_i_*]/2*N_e_* converges to 1 as *N_e_* becomes large.

### S3 Building pedigrees with Familinx

We simulated our Famlllnx-based pedigrees over `GEN` = 100 non-overlapping generations. For each generation, we selected family units at random from the data until the total number of children across all family units was greater than some pre-determined *N_c_* (the population census size). In addition, we required the total number of parents among the selected family units to be less than or equal to *N_c_*. Then, we connected the `GEN` generations together by randomly assigning each parent in generation *g* to be one of the children in generation *g* +1, disallowing sibling mating. Finally, we connected the first and last generation so that the pedigree is cyclical, with a period of `GEN` generations.

As a note, this procedure will not be appropriate for datasets where a substantial number of family units contain only one child, because the algorithm requires the number of children to be greater than or equal to the number of parents. When many families have only one child, families with more children will be over-sampled, and the family structure of our constructed pedigrees will be very different from the family structure we are attempting to replicate.

The value of *N_c_* was chosen to `GEN`erate pedigrees with a target effective population size, *N_e_*. For each *N_c_*, we estimated the effective population size of our pedigree by calculating the average time until coalescence over 50 sampled pairs, and setting *N_e_* as half of that time. We then discarded the pedigree unless this value is within *σ N_e_* of the target *N_e_*, where *σN_e_* is the standard deviation of the observed coalescent effective sizes for a population of size *N_c_* = *N_e_* in a Wright-Fisher model. We constrain our pedigrees to be close to the target effective population size because we want to make sure that the higher covariance we observe in the Familinx pedigree simulations relative to the WF model is not only due to potentially higher variance of *N_e_*.

We note that under our algorithm, some information on the country-specific pedigree structure is lost by breaking large genealogies into family units (e.g., inter-generational correlations in family size, or the rate of first and second cousin matings). Nevertheless, sufficient information is retained so that pedigrees with the same *N_e_* generated based on data from different countries are distinguished by their correlation of coalescence times (main text Figure 4).

